# The ABA receptor antagonist Antabactin restores germination of thermoinhibited lettuce

**DOI:** 10.1101/2024.02.16.580753

**Authors:** James Eckhardt, Sean Cutler

## Abstract

Germination is critical to agricultural productivity because low germination rates and/or asynchronous germination negatively affect stand establishment and subsequent yields. Exposure to high temperatures during seed imbibition can decrease both germination synchrony and rates through an ABA-mediated process called thermoinhibition. Methods to reduce thermoinhibition would be agriculturally valuable, particularly with increasing global mean temperatures. Lettuce seed germination is particularly sensitive to high temperatures and is a classic system for studying thermoinhibition. Extensive evidence using mutants and carotenoid biosynthetic inhibitors (*e.g*. fluridone) has demonstrated that endogenous abscisic acid (ABA) biosynthesis is required for thermoinhibition in lettuce and Arabidopsis. In principle, ABA biosynthetic inhibitors could be used to block thermoinhibition, but carotenoid biosynthetic inhibitors are not well-suited for this application due to their herbicidal effects and current hydroxamate ABA biosynthetic inhibitors lack potency. Here we explore the potential of ABA receptor antagonism to disrupt thermoinhibition using Antabactin (ANT), a broad spectrum high-affinity receptor antagonist. We show low µM ANT treatments (10µM) during seed imbibition reduce thermoinhibition at temperatures of up to 40°C, demonstrating that ABA signaling is required for thermoinhibition. We further explored interactions between ANT and seed priming, which is used commercially to improve seed germination and reduce thermoinhibition and is achieved by partial hydration and subsequent desiccation of seeds. We show that primed lettuce seeds have improved germination rates at high temperatures, consistent with previous observations; moreover, seeds primed in the presence of ANT show further increases in germination at elevated temperatures, showing that the two treatments can be combined to improve lettuce seed germination. Our data demonstrate that ABA antagonists are candidate agrochemicals for mitigating the effects of high temperature on seed germination and stand establishment that may be of increasing importance due to climate change.

## Introduction

In crop species, synchronous, rapid seed germination is necessary to optimize crop yield and quality. Seed dormancy is the failure of a viable dormant seed to germinate in conditions that are suitable for non-dormant seeds to germinate (Baskin and Baskin 2004). In wild plants, dormancy can be an advantageous trait. Among other adaptive advantages, it can optimize germination time seasonality and provide asynchronous germination, a bet hedging strategy (Burghardt, Edwards, and Donohue 2016; Simons and Johnston 2006). In crops however, seed dormancy, or lack thereof, can be a problem where controlled synchronous rapid germination is essential. Dormancy can lead to asynchronous emergence, causing issues in the homogenous stand establishment necessary for optimal management and dormancy can also impact the carefully controlled time farmers want emergence of crops to coincide with environmental conditions (Rodríguez et al. 2015). Dormancy is controlled in large part by the antagonism between two phytohormones ABA and GA (Gazzarrini and Tsai 2015; Graeber et al. 2012; Finkelstein et al. 2008). Additionally, DOG1 also plays an essential role as when either DOG1 or ABA is not present, seeds are not dormant (Bentsink et al. 2006; Nakabayashi et al. 2012; Kendall et al. 2011). A variety of environmental signals modulate ABA and GA levels, with temperature being highly important (Toh et al. 2008; Tamura et al. 2006).

When non-dormant seeds fail to germinate under high temperatures, this is called thermoinhibition. In the same way that dormancy and germination are controlled by antagonism between ABA and GA, the same is true for thermoinhibition. Thermoinhibition has been observed in multiple crops, however, lettuce is a model for thermoinhibition (Huo and Bradford 2015). Lettuce is a cool season crop, and in the United States over 95% of lettuce is produced in California and Arizona (“Vegetables 2022 Summary” 2023). As cropland is degraded and human related factors including, soil salinity, increasing temperatures and inhabitation of cropland, lettuce cultivation will become increasingly difficult. Since it is a cool season crop, lettuce germination is impacted by high temperatures, and being grown in California and Arizona, the seeds are often exposed to high temperatures during germination. Thus germination thermoinhibition, a failure to germinate at high temperatures, is a major problem in lettuce production causing reduced field emergence, stand establishment, and yield (Lafta and Mou 2013; Cantliffe, Guedes, and Shuler 1980). A variety of practices have been adopted to prevent crop loss to thermoinhibition: selection of heat tolerant varieties, seed priming and timing seed planting to occur at the coolest part of the day (Cantliffe, Guedes, and Shuler 1980; Holmes et al. 2019; Lafta and Mou 2013; Hill et al. 2006).

Due to the agronomic importance of thermoinhibition in lettuce, research has focused on understanding this process. The plant hormone Abscisic Acid (ABA) plays a major role in regulating germination thermoinhibition under high temperatures (Toh et al. 2008). Both Argyris et al 2008 and Gonai et al 2004 showed that endogenous levels of ABA in multiple lettuce varieties imbibed under high temperatures were higher than those of seeds imbibed at lower temperatures. In addition, the role of ABA in thermoinhibition was experimentally shown by both increased germination at high temperatures when fluridone, a carotenoid biosynthetic inhibitor was applied, and sensitivity to ABA increased under high temperatures (Jason Argyris et al. 2008; Gonai et al. 2004). Argyris et al 2008 also showed that sensitivity to heat in germination mapped to a QTL containing LsNCED4. Under high heat, lettuce seeds (*Lactuca sativa* ‘Salinas’) express LsNCED4 more highly and accumulate ABA to a greater extent (Jason Argyris et al. 2008). Taken together, these results suggest that ABA is an important regulator of germination thermoinhibition in Lettuce.

Recently Viadya et al. 2021 developed a potent synthetic ABA antagonist, Antabactin (ANT) which works by blocking the ABA receptor-PP2C interaction thereby reducing ABA signaling *in planta (Vaidya et al. 2021)*. ANT was shown to accelerate germination of Arabidopsis, tomato and barley. Additionally, ANT accelerated germination in thermoinhibited Arabidopsis. Since thermoinhibition of lettuce germination is known to be impacted by endogenous ABA, herein we utilize ANT to understand and regulate thermoinhibition. While previous research has used various indirect chemical and genetic methods to understand thermoinhibition, we have used ANT to directly block ABA signaling and can show the ABA contribution to thermoinhibition directly. We determine the potency ANT in thermoinhibited lettuce, compare it to common ABA biosynthetic inhibitors and determine the ABA effect on thermoinhibition at a variety of temperatures. Finally, we develop a seed priming method with ANT to improve germination rates at high temperatures.

## Methods

Seeds of *Lactuca sativa* ‘Salinas’ were grown in a greenhouse in summer 2021 and summer 2022 in Riverside, California. Plants were fully dried and seeds were collected and stored at room temperature for 3-5 months before experiments were carried out.

### Seed Germination Tests

#### Surface sterilization

Lettuce seeds were sterilized in room lighting and temperature using the following steps. Seeds were gently shaken for one minute in 70% ethanol. This was followed by fifteen minutes of gentle shaking in 20% bleach and three subsequent washes in sterile water. Seeds were immediately plated after sterilization.

#### Antabactin promotion of germination (EC50)

To establish the concentration of ANT at which 50% of seeds germinate under thermoinhibition, we conducted assays as follows. Surface sterilized Salinas seeds (n>35 seeds; seeds were harvested two months prior to the experiment) were plated in petri dishes on 0.7% agar medium containing ½ Murashige-Skoog (MS) and 10, 5, 1, 0.75, 0.5, 0.25, 0.1µM ANT and mock untreated control (0.1% v/v DMSO). Petri dishes were immediately transferred to 32°C and 25°C incubators in dark. After 48 hours, plates were photographed. Germination was scored as positive based on radicle emergence and the ET_50_ (time at which 50% of seeds have germinated) values were inferred by fitting the germination data over time to a 4 parameter log-logistic model in the drc (dose response curves) package (Ritz et al. 2015).

#### Chemical manipulation of ABA biosynthesis and signaling

To compare the effect of fluridone, AbamineSG and ANT on germination under thermoinhibition, we conducted the following assay. Surface sterilized Salinas seeds (n>35 seeds; seeds were harvested two months prior to the experiment) were plated in petri dishes on 0.7% agar mediµM containing 1/2 MS salts and either 10µM ANT, 30µM Fluridone, 100µM Fluridone, 30µM AbamineSG, 100µM AbamineSG or mock untreated control (1% v/v DMSO). Petri dishes were immediately transferred to 32°C and 25°C incubators in dark. After 48 hours, plates were photographed. Germination was scored upon radicle emergence.

#### Comparison with other hormone treatments

Surface sterilized Salinas seeds (n=40 seeds) were plated in petri dishes on 0.7% agar medium containing 1/2 MS salts and either 10µM ANT, 100µM GA, 100nM KAR_1_ or mock untreated control (1% v/v DMSO). Petri dishes were immediately transferred to 32°C and 25°C incubators in dark. After 48 hours, plates were photographed. Germination was scored upon radicle emergence.

#### Effective temperature range of ANT

To determine the upper temperature range at which ANT promotes germination we performed the following assay. Surface sterilized Salinas seeds (n>35 seeds; seeds were harvested two months prior to the experiment) were plated in petri dishes on 0.7% agar medium containing 1/2 MS salts and either 10µM ANT or mock untreated control (0.1% v/v DMSO). Petri dishes were immediately transferred to 25°C, 32°C, 37°C, 40°C and 45°C incubators in dark. Petri dishes were imaged at 48 hours and 120 hours. After the 120 hour time point, all plates were transferred to the 25°C incubator and imaged after 72 hours in dark at 25°C to determine if seeds were thermoinhibited or thermodormant. Seeds incubated at 45°C were stained with tetrazolium to determine viability. Staining was carried out as described in HCRM Catao et al. 2018 (Catão et al. 2018).

### Gene Expression Assays

For gene expression assays, two tissue types were used. Salinas seeds were grown on petri dishes on 0.7% agar medium containing 1/2 MS salts and either 10µM ANT or mock untreated control (0.1% v/v DMSO). Seeds were grown for 24 hours in dark before RNA extraction. Each treatment was conducted in biological triplicate. 50mg of tissue was collected and used for RNA extraction. RNA was also extracted from Salinas leaf tissue. 100mg of leaf tissue was cut and placed in petri dishes with either 100µM ABA, 100µM ABA and 50µM ANT or mock untreated control (0.1% v/v DMSO) in 10mL water. Leaf tissue was incubated at 25°C for 8 hours in 70µM light before RNA extraction. Each treatment was conducted in biological triplicate. The total RNA was extracted using the RNeasy Plant Mini Kit (QIAGEN, Cat No. 74904), and cDNA was synthesized by QuantiTect Reverse Transcription Kit (QIAGEN, Cat No. 205311). Quantitative RT-PCR was performed with CFX Connect Real-Time PCR Detection System using Maxima SYBR Green/Fluorescein qPCR Master Mix (Thermo Scientific, Cat No. K0242). The relative expression level of the genes was normalized against the reference gene UBC21 by ΔCT Method. Statistical tests were performed using the raw ΔCT values. A pairwise t test was used to assess the differences between treatments.

The primers used in the experiments are listed as follows.

LsUBC21: 5’TCTTAGATCACCGTCCCATCGT3’, 5’TCTGAGATTGTCCGAGGATATGAG3’

LsNCED4: 5’TGATCCAGCGGTTCAGCTAA3’, 5’TTCACCAATTACCTCCAGACCAT3’

LsM3K18: 5’ATGATTGAAATGGCCACCGG3’, 5’TGATCGGAAAATTCATCTGGGA3’

### Seed Priming

Seeds were primed in -1.25mPa PEG8000 (0.2968g/mL). Water potential was calculated as in Michel 1983 (Michel 1983). Seeds were placed in petri dishes on blotting paper with 10mL of PEG solution and either 10µM ANT or mock untreated control (0.1% v/v DMSO). Seeds were at 9°C in low light (1µM) for 48hrs, rinsed three times with water, dried at 32°C in an incubator for 2 hours and then stored for >48 hours at 9°C in dark before use. For germination experiments, seeds were surface sterilized and plated in petri dishes on 0.7% agar medium containing 1/2 MS salts. Seeds were incubated in the dark at 35°C for 24 hours before germination was quantified.

Approximately 10mg of primed seeds were extracted in 500uL MeOH. Seeds were put through two homogenization cycles in a Precellys 24. Three 10 second cycles of 6500rpm were used. After homogenization, tubes were centrifuged at 13.3rpm for 2 minutes and 100uL of supernatant was used for LCMS. For ANT spiked samples, a 1% v/v solution of ANT in MeOH was added to PEG only primed seeds to make 125, 250, 500 and 1000nM final concentrations. The analysis of ANT was performed on an Agilent 1260 HPLC system coupled with an Agilent 6224 ESI TOF in positive ion mode. Chromatographic separation was carried out on an Agilent Poroshell 120 EC-C18 column (50 x 3mm, 2.7µM) using mobile phases A (water/formic acid, 100/0.1, v/v) and B (acetonitrile/formic acid, 100/0.1, v/v). The flow rate was 0.5mL/min and the injection volume was 5uL.

### Phylogeny

The coding regions of the 21 mitogen activated protein kinase kinase kinases (MAP3K) were downloaded from TAIR. Homologous genes to MAP3K18 in Lactuca sativa were identified by using NCBI blastn, and three homologous sequences were selected. Using MEGA (Version 11.0.13) the nucleotide sequences were aligned by MUSCLE (Tamura K, Stecher G, and Kumar S 2021). A maximum likelihood tree was then constructed in MEGA using default settings.

## Results

ANT restores germination in thermoinhibited plants as effectively as fluridone. At 32°C in dark conditions, *L. sativa* ‘Salinas’ germination was greatly reduced, with mean germination between 0-18% after two days (Fig. 1A). At 25°C, seeds showed 100% germination throughout the experiments conducted, and fluridone (10µM), a carotenoid biosynthetic inhibitor fully restored germination to 100%. Additionally, ANT treatment (10µM) also nearly fully restored germination under high temperatures (97% germination). AbamineSG, an NCED inhibitor, did not promote germination. In addition to chemical inhibitors, ANT was compared to other exogenous and endogenous germination related chemicals. ANT is more potent than gibberellic acid (GA) karrikin 1 (KAR_1_) (Fig. 1B) treatments. GA (100µM) and KAR_1_ (100nM) treatments led to similar germination rates to mock after two days in dark at 32°C, 8%, 10% and 18% respectively. As before, 10µM ANT nearly fully restored germination (94%).

**Figure 1.**
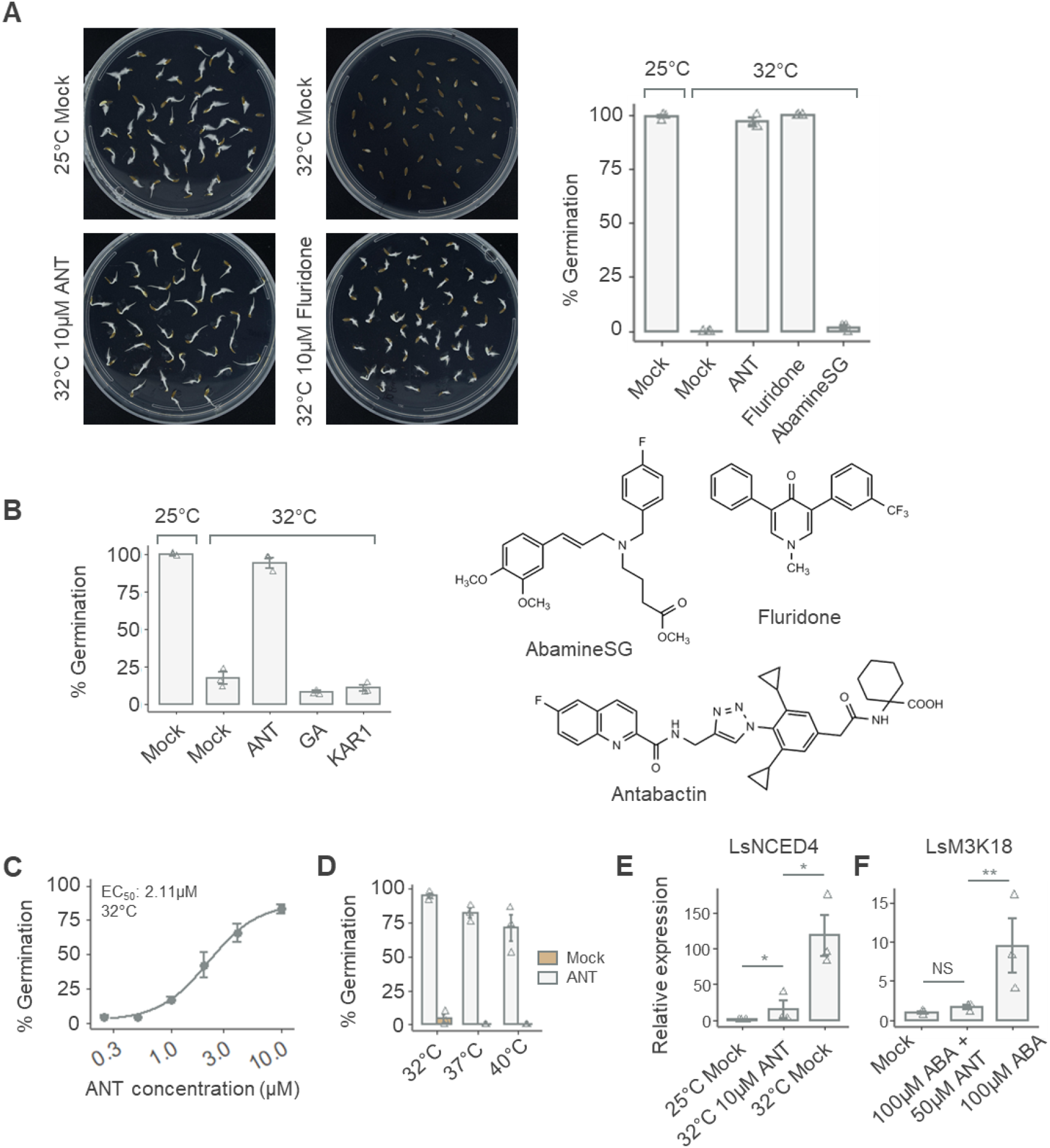
(A) ANT and fluridone reverse thermoinhibition in L. sativa (‘Salinas’). Germination was quantified for seeds imbibed on plates containing DMSO (mock treated), 10µM ANT, 10µM fluridone or 100µM AbamineSG and grown for 48h. (B) GA and KAR1 did not reverse thermoinhibition. Germination was quantified for seeds imbibed on plates contained DMSO, 10µM ANT, 100µM GA or 100nM KAR1 and grown for 48h. (C) ANT is potent in the low micromolar range. Germination was quantified for seeds after growth on plates for 24h at 32°C. ET_50_ value inset in graph are inferred by nonlinear fits in drc. (D) 10µM ANT reverses thermoinhibition at temperatures of up to 40°C. Germination was quantified seeds after growth in either ANT or mock (DMSO) for 48h. (E) ANT treatment reduces NCED4 expression (normalized to UBC21) measured by qRT-PCR of seeds grown for 24 hours in mock (DMSO) or 10µM ANT. (F) ANT treatment reduces the M3K18 related expression (normalized to UBC21) measured by qRT-PCR of lettuce leaves in (DMSO) or 100µM ABA or 100µM ABA and 50µM ANT in water. * indicates p < 0.05, ** indicates p < 0.01 (pairwise two sample t test corrected for multiple comparisons). In (A), (B), (C), (D) and (E) liquid sterilized seeds were grown on ½ MS, 0.7% agar plates in dark. Error bars indicate SEM (n=3).

ANT is potent in the low micromolar range and at high temperatures. The EC_50_ of ANT is 2.11µM (Fig. 1C). Lettuce seeds were grown for two days in dark at 32°C in six ANT concentrations (0.25, 0.5, 1, 2.5, 5 and 10µM). 10µM ANT nearly fully restored germination (91% mean) and 0.25µM ANT had little effect (7% mean). Since 10µM ANT restored germination under thermoinhibition, we tested to determine the upper temperature limit of ANT effect. After 120hrs, ANT promoted germination at 37°C and 40°C (Fig. 1D). At 37°C, 10µM ANT treatment resulted in 82% germination and at 40°C ANT treatment resulted in 74% germination. Germination was also quantified at 48 hours, and the higher temperatures slowed germination at 37°C and 40°C leading to only 47% and 8% germination respectively (Fig. S4). At 45°C lettuce seeds were dead and no longer viable (Fig. S2). Seeds at 45°C looked visibly unviable at 120 hours and this was confirmed through tetrazolium staining.

ANT reduces expression of ABA and thermoinhibition upregulated genes. LsNCED4, a homolog to AtNCED6, is upregulated under thermoinhibition. At 32°C in dark after 24 hours, it was upregulated 118-fold while at the same temperature NCED4 in ANT treated seeds was only upregulated 15-fold over mock at 25°C (Fig. 1E). After 24 hours the mock and ANT treated seeds begin to germinate, while the mock seeds will likely not germinate. The seeds are thus in different developmental stages. By applying ANT, ABA and a combination of the two molecules to leaf tissue we eliminated any developmental effects and showed that while LsNCED4 is not upregulated by endogenous ABA (Fig. S3), a homolog (Lsat_1_v5_gn_7_91420) to AtMAPKKK18 is significantly upregulated (9.5 fold increase) under endogenous ABA treatment (100µM) and downregulated by a coapplication of ANT (50µM) and ABA (100µM) (Fig. 1F and Fig. S1).

Osmotic priming with ANT leads to ANT uptake and storage within lettuce seeds. Three different batches of seeds primed in a solution of 10µM ANT and -1.25mPa PEG showed they contained ANT (Fig. 2B). Seeds were primed, dried and stored for 5-7 days before extraction and ANT level quantification and all showed ANT similar to seeds primed with only PEG and later spiked with 500nM ANT before LCMS analysis. The three primed seed batches showed an average of 30% higher germination rates when primed with ANT than solely PEG at 35°C after 24 hours (Fig. 2A). The seeds that were after ripened the longest had the highest germination under both ANT and mock, 90% and 52% respectively seeds with the shortest after ripening, 1.5 months, had the lowest germination 30% and 9%.

**Figure 2.**
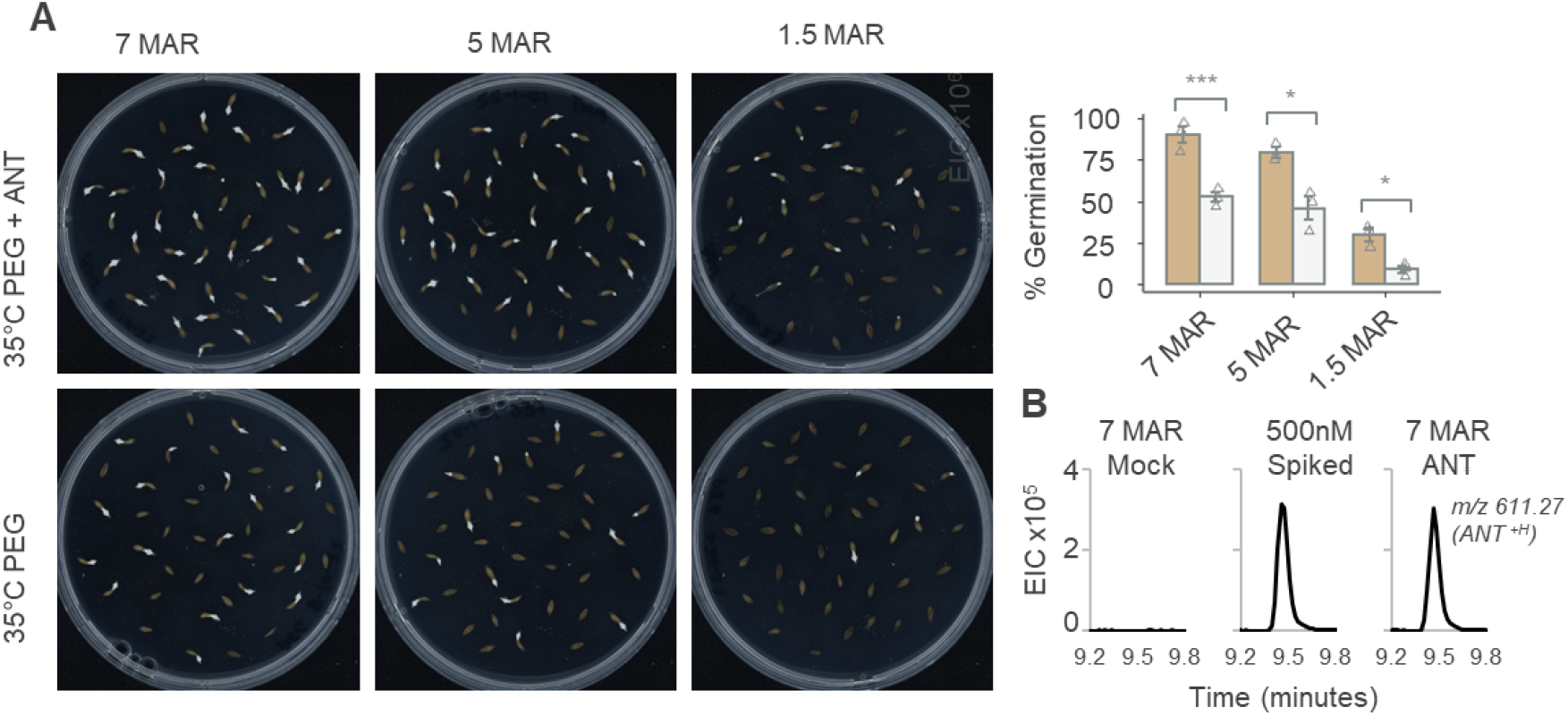
Priming seeds with ANT improves germination rates under thermoinhibition. (A) Germination was quantified for seeds primed in -1.25 Mpa solution of PEG 8000 and either 100 µM ANT or mock (DMSO). Primed seeds were plated on ½ MS, 0.7% agar plates and germination was quantified at 24h in 35°C. *** indicates p < 0.01 (two sample t test). Error bars indicate SEM (n=3). (B) Extracted ion chromatograms (EIC) for ANT^+H^ m/z 611.270 (+/-50 ppm).

## Discussion

In wild plants, dormancy is an adaptive trait which has direct impacts on fitness. Timing of germination is important so plants can start their life cycle at the beginning of a growth season in optimal conditions. Seeds from a single parent can also show variation in dormancy, a bet hedging strategy, which reduces risk of the mortality of all progeny. While dormancy can be advantageous in the wild, the trait is not beneficial in crops where synchronous, fast emergence is necessary to maximize optimal environmental conditions (Rodríguez et al. 2015). As such, low dormancy has been selected in many crops, however various environmental factors can still induce dormancy. The phytohormone abscisic acid (ABA) is widely known to be one important dormancy regulator in seeds and has previously been implicated in thermoinhibition (Finch-Savage and Leubner-Metzger 2006; Gonai et al. 2004; Toh et al. 2008). Lettuce seeds are thermoinhibited at temperatures below the limit for healthy seedling growth, and ABA is an important regulator of this thermoinhibition (Cantliffe, Shuler, and Gliedes 1981; Gonai et al. 2004). Exogenous ABA can reduce the upper limit for germination and upon thermoinhibition, endogenous ABA levels increase (Jason Argyris et al. 2008).

Herein we use a new tool to validate ABA’s regulation of thermoinhibition by directly blocking ABA signaling with the ABA antagonist Antabactin. I will discuss the chemical manipulations of ABA first and then discuss the genetic findings in thermoinhibition. To understand the extent of ABA’s regulation of germination, we can first compare it to other chemicals known to influence germination. The phytohormone GA promotes seed germination. In part it does this through negatively affecting ABA biosynthesis and signaling and promoting ABA catabolism. Argyris et al 2008 showed that the GA effect of germination in lettuce seeds was variety dependent, with *L. sativa* ‘Salinas’ being less affected by GA application (Jason Argyris et al. 2008). This held true in my experiments with GA not significantly increasing germination rates over mock at 32°C. Karrikins have also been shown to improve germination in lettuce (Martinez et al. 2022). Karrikins are chemicals in smoke that can enhance germination, and in *L. sativa* ‘Grand Rapids’, they have improved germination rates under adverse conditions. Again, in *L. sativa* ‘Salinas’, KAR_1_ did not improve germination rates under mock under temperatures inducing thermoinhibition. This could be a variety based issue or suggests thermoinhibition is not influenced by karrikins. Fluridone has previously been shown to improve germination under thermoinhibition by reducing *de novo* ABA synthesis (Yoshioka, Endo, and Satoh 1998; Roth-Bejerano et al. 1999). Fluridone is a carotenoid biosynthetic inhibitor. It acts by inhibiting phytoene desaturase which converts phytoene to phytofluene, an early precursor of ABA (Bartels and Watson 1978; Fong and Schiff 1979). In many species, fluridone promotes and accelerates germination under thermoinhibition. This is also true of lettuce, where application of fluridone promotes germination and reduces the ABA content in imbibed seeds (Yoshioka, Endo, and Satoh 1998). Fluridone application results in reduced *de novo* ABA biosynthesis, but is an imperfect inhibitor as it also reduces synthesis of many other carotenoids which are often signaling molecules in plants, including strigolactone, another phytohormone. Argyris et al showed that 100µM fluridone application restored germination in dark at 32°C, and Fig 1A shows that this is the case (Jason Argyris et al. 2008). While fluridone has some application as an inhibitor, its herbicidal effects give it no agricultural application in this regard. Taken together, this suggests that ABA is the primary signal for thermoinhibition in *L. sativa* ‘Salinas’. All this evidence is, however, indirect. We directly confirmed these conclusions by utilizing the chemical tool ANT. We applied ANT to thermoinhibited lettuce seeds, and it fully restored germination. ANT is a ABA antagonist, competitively inhibiting ABA binding to the PYR/PYL ABA receptors, and the germination response is that of if ABA signaling was blocked. ANT blocks signaling from both new ABA biosynthesis and existing ABA pools. While previous approaches have genetically or chemically indirectly altered ABA signaling, we block ABA signaling directly leading to restored germination and this confirms that ABA is the primary regulator of thermoinhibition in lettuce. We previously showed that ANT also reduces time to germination in heat treated A. thaliana seeds, directly demonstrating ABA regulates thermoinhibition in multiple species and that ANT is a valuable tool to assess this across species (Vaidya et al. 2021). Previous research has also suggested that the NCEDs are the rate limiting step in ABA biosynthesis. Abamine is an NCED inhibitor and reduces ABA accumulation in planta and promotes seed germination, however, it is phytotoxic (Han, Kitahata, Sekimata, et al. 2004; Han, Kitahata, Saito, et al. 2004). More recently, AbamineSG was developed and demonstrated reduced phytoxicity and led to reduced ABA content by NCED inhibition (Schwartz, Qin, and Zeevaart 2003; Kitahata et al. 2006). In our experiments, AbamineSG application did not improve germination under thermoinhibition, likely due to ineffective action rather than the NCEDs not being important.

To further confirm ANT’s action in block ABA signaling, we looked at mRNA levels of ABA and thermoinhibition regulated genes. ANT reduces thermoinhibition by blocking ABA signaling and likely influencing ABA regulated gene expression. In Arabidopsis, ANT restores transcript levels of classic ABA upregulated genes, and we wanted to confirm this action in lettuce. M3K18 is an ABA marker gene in Arabidopsis, being highly upregulated under ABA treatment or osmotic stress. ANT treatment restores transcript levels to those of mock. M3K18 is closely related in sequence and trasncriptional profile to M3K17. These two genes are functionally redundant (Danquah et al. 2015). Three lettuce homologs cluster with the two genes. We chose one, 91420 to test as an ABA marker gene in lettuce. In lettuce, exogenous ABA application led to increased expression in leaf disks, and ANT reduced expression so they were not significantly different from mock, suggesting that ANT is acting through restoring normal levels of expression to ABA regulated genes. This suggests that 91420 is a good candidate for an ABA marker gene for future studies in lettuce. LsNCED4 is a well known factor in influencing thermoinhibition in lettuce (Jason Argyris et al. 2008; Bertier et al. 2018; Huo et al. 2013). LsNCED4 is the homolog of AtNCED6 (Sawada et al. 2008). NCED catalyzes a rate limiting step in ABA biosynthesis, the oxidative cleavage of 9-cis vioxanthin or neoxanthin to xanthoxin, and thus loss of function of the NCEDs results in reduced ABA biosynthesis. This is also the first committed step in ABA biosynthesis (Schwartz, Qin, and Zeevaart 2003; Nambara and Marion-Poll 2005). Expression of LsNCED4 was previously shown to increase during thermoinhibition, and Fig 1E also confirms this (Jason Argyris et al. 2008). Additionally, deletions in LsNCED4 were shown to dramatically increase the maximum temperature at which lettuce could germinate (Bertier et al. 2018). This directly implicates ABA biosynthesis and thus levels and signaling lead to thermoinhibition. ANT application, which blocks ABA signaling reduces expression of NCED4, however this effect may not be direct as seeds imbibed for 24 hours in both mock and ANT treatments are beginning to germination and mock at 32°C are not germination. They are in different developmental stages which could mean that NCED4 expression naturally decreases as seeds germinate. Both M3K18 and NCED4 expression patterns further demonstrate ANT’s action is a result of transcriptional regulation of ABA and thermoinhibition regulated genes.

These results suggest ANT’s action through reducing ABA signaling. When ABA signaling is blocked by ANT in Arabidopsis, barley and tomato, we previously showed time to germination decreases (Vaidya et al. 2021). In lettuce under thermoinhibition, there is little to no germination, and blocking ABA signaling with ANT fully restores germination. We calculated an EC50 for ANT at 2.11µM at 32C, and, at 10µM concentrations, germination is fully restored. As such, we used 10µM ANT for further germination tests. Argyris et al. 2008 showed that germination of *L. sativa* ‘Salinas’ decreases at 27°C with germination being fully thermoinhibited at 29°C (J. Argyris et al. 2008). Thermoinhibition is present across many additional crisphead lettuce varieties with Lafta and Mou 2013 showing that germination of all crisphead lettuce varieties tested at 34°C, none had higher than 80% germination with most not germinating above 50% (Lafta and Mou 2013). We wanted to determine the extent to which ABA contributes to thermoinhibition even at very high temperatures. Blocking ABA signaling at temperatures of 40°C results in 74% germination. This shows that ABA is the major contributor to thermoinhibition in lettuce even at very high temperatures. However, upon restoration to normal temperatures, 100% of seeds germinated, suggesting one of two things. Either the ABA effect is not the only factor contributing to thermoinhibition or ANT is not fully blocking ABA signaling. Of note is that germination did slow under 37°C and 40°C, and while at 32°C nearly 100% of seeds were germinated at 48 hours, it took 120 hours before 40°C germination reached 74%. Temperatures of 45°C killed seeds, also suggesting that ABA is the major contributor to thermoinhibition up to the point where seeds are no longer viable.

In agriculture, final emergence rate and synchrony of germination are important factors in determining final crop yield, and, as ABA is the major contributor to thermoinhibition, we wanted to determine if there could be an agriculturally relevant way to apply ANT to improve seed emergence. Previously, targeting NCEDs through Crispr has been discussed as a way to target ABA induced thermoinhibition, and it is successful in restoring germination, however this method is not without disadvantages (Bertier et al. 2018). Firstly, gene editing is regulated, and this process of editing must be done to every variety of lettuce grown to produce the desired effect. Secondly, lettuce is grown in a hot climate, and the NCEDs are important parts of the ABA biosynthetic pathway. ABA controls stomatal aperture and thus transpiration, and altering these processes may be harmful to lettuce plants later in their growth (Cutler et al. 2010). Pre germination seed treatments can mitigate these issues. As such, we osmotically primed lettuce seeds in PEG and ANT to improve their germination under thermoinhibition. Osmotic priming is a method by which seeds are rehydrated to a set osmotic potential under which seeds begin to germinate but radicle protrusion cannot occur. Seeds are subsequently redried before later planting. Priming has already been shown to improve germination traits (Cantliffe, Shuler, and Gliedes 1981; Bradford 1986; Paparella et al. 2015). Additionally, hormonal priming with ABA reduces germination suggesting that a hormonal priming treatment targeting the ABA pathway can have lasting effects (Tarquis and Bradford 1992). Our priming treatment resulted in ANT uptake in the seeds, as shown by LCMS. Spiking seeds with 500nM ANT showed the same peak area as the primed seeds, suggesting that the concentration of ANT in primed seeds was at least somewhat similar to the 500nM concentration. Priming with ANT and PEG over just PEG promoted higher germination rates, and could be potentially beneficial in a agricultural setting by greatly reducing ABA signaling. ANT priming has potential to be applied to other crops where ABA signaling blocks germination. In light of increased global food demand and the variety of abiotic and biotic factors challenging food production, crop production will need to expand to less suitable growing environments. As this production expands, so will the challenges facing growers. ANT provides another tool to meet these growing challenges.

## Notes

### Competing Interest Statement

The authors have declared no competing interest.

## References

Argyris, Jason, Peetambar Dahal, Eiji Hayashi, David W. Still, and Kent J. Bradford. 2008. “Genetic Variation for Lettuce Seed Thermoinhibition Is Associated with Temperature-Sensitive Expression of Abscisic Acid, Gibberellin, and Ethylene Biosynthesis, Metabolism, and Response Genes.” Plant Physiology 148 (2): 926–47.

Argyris, J., P. Dahal, M. J. Truco, O. Ochoa, D. W. Still, R. W. Michelmore, and K. J. Bradford. 2008. “Genetic Analysis of Lettuce Seed Thermoinhibition.” Acta Horticulturae, no. 782 (February): 23–34.

Bartels, P. G., and C. W. Watson. 1978. “Inhibition of Carotenoid Synthesis by Fluridone and Norflurazon.” Weed Science 26 (2): 198–203.

Baskin, Jerry M., and Carol C. Baskin. 2004. “A Classification System for Seed Dormancy.” Seed Science Research 14 (1): 1–16.

Bentsink, Leónie, Jemma Jowett, Corrie J. Hanhart, and Maarten Koornneef. 2006. “Cloning of DOG1, a Quantitative Trait Locus Controlling Seed Dormancy in Arabidopsis.” Proceedings of the National Academy of Sciences of the United States of America 103 (45): 17042–47.

Bertier, Lien D., Mily Ron, Heqiang Huo, Kent J. Bradford, Anne B. Britt, and Richard W. Michelmore. 2018. “High-Resolution Analysis of the Efficiency, Heritability, and Editing Outcomes of CRISPR/Cas9-Induced Modifications of NCED4 in Lettuce (Lactuca Sativa).” G3 8 (5): 1513–21.

Bradford, Kent J. 1986. “Manipulation of Seed Water Relations via Osmotic Priming to Improve Germination under Stress.” HortScience: A Publication of the American Society for Horticultural Science 21 (5): 1105–12.

Burghardt, Liana T., Brianne R. Edwards, and Kathleen Donohue. 2016. “Multiple Paths to Similar Germination Behavior in Arabidopsis Thaliana.” The New Phytologist 209 (3): 1301–12.

Cantliffe, D. J., A. C. Guedes, and K. D. Shuler. 1980. “OVERCOMING THERMODORMANCY IN A HEAT-SENSITIVE ROMAINE LETTUCE SEED BY PRIMING.” In HORTSCIENCE, 15:405–405. AMER SOC HORTICULTURAL SCIENCE 701 NORTH SAINT ASAPH STREET, ALEXANDRIA, VA ….

Cantliffe, D. J., K. D. Shuler, and A. C. Gliedes. 1981. “Overcoming Seed Thermodormancy in a Heat Sensitive Romaine Lettuce by Seed Priming1.” HortScience: A Publication of the American Society for Horticultural Science 16 (2): 196–98.

Catão, Hugo Cesar R. M., Luiz Antonio Augusto Gomes, Renato M. Guimarães, Pedro Henrique F. Fonseca, Franciele Caixeta, and Alexandre G. Galvão. 2018. “Physiological and Biochemical Changes in Lettuce Seeds during Storage at Different Temperatures.” Horticultura Brasileira 36 (1): 118–25.

Cutler, Sean R., Pedro L. Rodriguez, Ruth R. Finkelstein, and Suzanne R. Abrams. 2010. “Abscisic Acid: Emergence of a Core Signaling Network.” Annual Review of Plant Biology 61: 651–79.

Danquah, Agyemang, Axel de Zélicourt, Marie Boudsocq, Jorinde Neubauer, Nicolas Frei Dit Frey, Nathalie Leonhardt, Stephanie Pateyron, et al. 2015. “Identification and Characterization of an ABA-Activated MAP Kinase Cascade in Arabidopsis Thaliana.” The Plant Journal: For Cell and Molecular Biology 82 (2): 232–44.

Finch-Savage, William E., and Gerhard Leubner-Metzger. 2006. “Seed Dormancy and the Control of Germination.” The New Phytologist 171 (3): 501–23.

Finkelstein, Ruth, Wendy Reeves, Tohru Ariizumi, and Camille Steber. 2008. “Molecular Aspects of Seed Dormancy.” Annual Review of Plant Biology 59: 387–415.

Fong, F., and J. A. Schiff. 1979. “Blue-Light-Induced Absorbance Changes Associated with Carotenoids in Euglena.” Planta 146 (2): 119–27.

Gazzarrini, Sonia, and Allen Yi-Lun Tsai. 2015. “Hormone Cross-Talk during Seed Germination.” Essays in Biochemistry 58: 151–64.

Gonai, Takeru, Shusuke Kawahara, Makoto Tougou, Shigeru Satoh, Teruyoshi Hashiba, Nobuhiro Hirai, Hiroshi Kawaide, Yuji Kamiya, and Toshihito Yoshioka. 2004. “Abscisic Acid in the Thermoinhibition of Lettuce Seed Germination and Enhancement of Its Catabolism by Gibberellin.” Journal of Experimental Botany 55 (394): 111–18.

Graeber, Kai, Kazumi Nakabayashi, Emma Miatton, Gerhard Leubner-Metzger, and Wim J. J. Soppe. 2012. “Molecular Mechanisms of Seed Dormancy.” Plant, Cell & Environment 35 (10): 1769–86.

Han, Sun-Young, Nobutaka Kitahata, Tamio Saito, Masatomo Kobayashi, Kazuo Shinozaki, Shigeo Yoshida, and Tadao Asami. 2004. “A New Lead Compound for Abscisic Acid Biosynthesis Inhibitors Targeting 9-Cis-Epoxycarotenoid Dioxygenase.” Bioorganic & Medicinal Chemistry Letters 14 (12): 3033–36.

Han, Sun-Young, Nobutaka Kitahata, Katsuhiko Sekimata, Tamio Saito, Masatomo Kobayashi, Kazuo Nakashima, Kazuko Yamaguchi-Shinozaki, Kazuo Shinozaki, Shigeo Yoshida, and Tadao Asami. 2004. “A Novel Inhibitor of 9-Cis-Epoxycarotenoid Dioxygenase in Abscisic Acid Biosynthesis in Higher Plants.” Plant Physiology 135 (3): 1574–82.

Hill, Hank, Kent J. Bradford, Jesse Cunningham, and Alan G. Taylor. 2006. “Primed Lettuce Seeds Exhibit Increased Sensitivity to Moisture during Aging.” In IV International Symposium on Seed, Transplant and Stand Establishment of Horticultural Crops; Translating Seed and Seedling 782, 135–42. actahort.org.

Holmes, Sydney C., Daniel E. Wells, Jeremy M. Pickens, and Joseph M. Kemble. 2019. “Selection of Heat Tolerant Lettuce (Lactuca Sativa L.) Cultivars Grown in Deep Water Culture and Their Marketability.” Horticulturae 5 (3): 50.

Huo, Heqiang, and Kent J. Bradford. 2015. “Molecular and Hormonal Regulation of Thermoinhibition of Seed Germination.” In Advances in Plant Dormancy, edited by James V. Anderson, 3–33. Cham: Springer International Publishing.

Huo, Heqiang, Peetambar Dahal, Keshavulu Kunusoth, Claire M. McCallum, and Kent J. Bradford. 2013. “Expression of 9-Cis-EPOXYCAROTENOID DIOXYGENASE4 Is Essential for Thermoinhibition of Lettuce Seed Germination but Not for Seed Development or Stress Tolerance.” The Plant Cell 25 (3): 884–900.

Kendall, Sarah L., Anja Hellwege, Poppy Marriot, Celina Whalley, Ian A. Graham, and Steven Penfield. 2011. “Induction of Dormancy in Arabidopsis Summer Annuals Requires Parallel Regulation of DOG1 and Hormone Metabolism by Low Temperature and CBF Transcription Factors.” The Plant Cell 23 (7): 2568–80.

Kitahata, Nobutaka, Sun-Young Han, Natsumi Noji, Tamio Saito, Masatomo Kobayashi, Takeshi Nakano, Kazuyuki Kuchitsu, et al. 2006. “A 9-Cis-Epoxycarotenoid Dioxygenase Inhibitor for Use in the Elucidation of Abscisic Acid Action Mechanisms.” Bioorganic & Medicinal Chemistry 14 (16): 5555–61.

Lafta, Abbas, and Beiquan Mou. 2013. “Evaluation of Lettuce Genotypes for Seed Thermotolerance.” HortScience: A Publication of the American Society for Horticultural Science 48 (6): 708–14.

Martinez, Stephanie E., Caitlin E. Conn, Angelica M. Guercio, Claudia Sepulveda, Christopher J. Fiscus, Daniel Koenig, Nitzan Shabek, and David C. Nelson. 2022. “A KARRIKIN INSENSITIVE2 Paralog in Lettuce Mediates Highly Sensitive Germination Responses to Karrikinolide.” Plant Physiology 190 (2): 1440–56.

Michel, B. E. 1983. “Evaluation of the Water Potentials of Solutions of Polyethylene Glycol 8000 Both in the Absence and Presence of Other Solutes.” Plant Physiology 72 (1): 66–70.

Nakabayashi, Kazumi, Melanie Bartsch, Yong Xiang, Emma Miatton, Silke Pellengahr, Ryoichi Yano, Mitsunori Seo, and Wim J. J. Soppe. 2012. “The Time Required for Dormancy Release in Arabidopsis Is Determined by DELAY OF GERMINATION1 Protein Levels in Freshly Harvested Seeds.” The Plant Cell 24 (7): 2826–38.

Nambara, Eiji, and Annie Marion-Poll. 2005. “Abscisic Acid Biosynthesis and Catabolism.” Annual Review of Plant Biology 56: 165–85.

Paparella, S., S. S. Araújo, G. Rossi, M. Wijayasinghe, D. Carbonera, and Alma Balestrazzi. 2015. “Seed Priming: State of the Art and New Perspectives.” Plant Cell Reports 34 (8): 1281–93.

Ritz, Christian, Florent Baty, Jens C. Streibig, and Daniel Gerhard. 2015. “Dose-Response Analysis Using R.” PloS One 10 (12): e0146021.

Rodríguez, María V., José M. Barrero, Francoise Corbineau, Frank Gubler, and Roberto L. Benech-Arnold. 2015. “Dormancy in Cereals (not Too Much, Not so Little): About the Mechanisms behind This Trait.” Seed Science Research 25 (2): 99–119.

Roth-Bejerano, Nurit, Norbert J. A. Sedee, Rene M. van der Meulen, and Mei Wang. 1999. “The Role of Abscisic Acid in Germination of Light-Sensitive and Light-Insensitive Lettuce Seeds.” Seed Science Research 9 (2): 129–34.

Sawada, Yoshiaki, Miki Aoki, Kentaro Nakaminami, Wataru Mitsuhashi, Kiyoshi Tatematsu, Tetsuo Kushiro, Tomokazu Koshiba, et al. 2008. “Phytochrome- and Gibberellin-Mediated Regulation of Abscisic Acid Metabolism during Germination of Photoblastic Lettuce Seeds.” Plant Physiology 146 (3): 1386–96.

Schwartz, Steven H., Xiaoqiong Qin, and Jan A. D. Zeevaart. 2003. “Elucidation of the Indirect Pathway of Abscisic Acid Biosynthesis by Mutants, Genes, and Enzymes.” Plant Physiology 131 (4): 1591–1601.

Simons, Andrew M., and Mark O. Johnston. 2006. “Environmental and Genetic Sources of Diversification in the Timing of Seed Germination: Implications for the Evolution of Bet Hedging.” Evolution; International Journal of Organic Evolution 60 (11): 2280–92.

Tamura K, Stecher G, and Kumar S. 2021. MEGA11: Molecular Evolutionary Genetics Analysis (version Version 11). http://www.megasoftware.net.

Tamura, Noriko, Takahiro Yoshida, Arata Tanaka, Ryuta Sasaki, Asuka Bando, Shigeo Toh, Loïc Lepiniec, and Naoto Kawakami. 2006. “Isolation and Characterization of High Temperature-Resistant Germination Mutants of Arabidopsis Thaliana.” Plant & Cell Physiology 47 (8): 1081–94.

Tarquis, Ana M., and Kent J. Bradford. 1992. “Prehydration and Priming Treatments That Advance Germination Also Increase the Rate of Deterioration of Lettuce Seeds.” Journal of Experimental Botany 43 (3): 307–17.

Toh, Shigeo, Akane Imamura, Asuka Watanabe, Kazumi Nakabayashi, Masanori Okamoto, Yusuke Jikumaru, Atsushi Hanada, et al. 2008. “High Temperature-Induced Abscisic Acid Biosynthesis and Its Role in the Inhibition of Gibberellin Action in Arabidopsis Seeds.” Plant Physiology 146 (3): 1368–85.

Vaidya, Aditya S., Francis C. Peterson, James Eckhardt, Zenan Xing, Sang-Youl Park, Wim Dejonghe, Jun Takeuchi, et al. 2021. “Click-to-Lead Design of a Picomolar ABA Receptor Antagonist with Potent Activity in Vivo.” Proceedings of the National Academy of Sciences of the United States of America 118 (38). 10.1073/pnas.2108281118.

“Vegetables 2022 Summary.” 2023. United States Department of Agriculture.

Yoshioka, Toshihito, Takashi Endo, and Shigeru Satoh. 1998. “Restoration of Seed Germination at Supraoptimal Temperatures by Fluridone, an Inhibitor of Abscisic Acid Biosynthesis.” Plant & Cell Physiology 39 (3): 307–12.

