## SupplementalFigures for "The ABA receptor antagonist Antabactin restores germination of thermoinhibited lettuce"

**Supplemental**

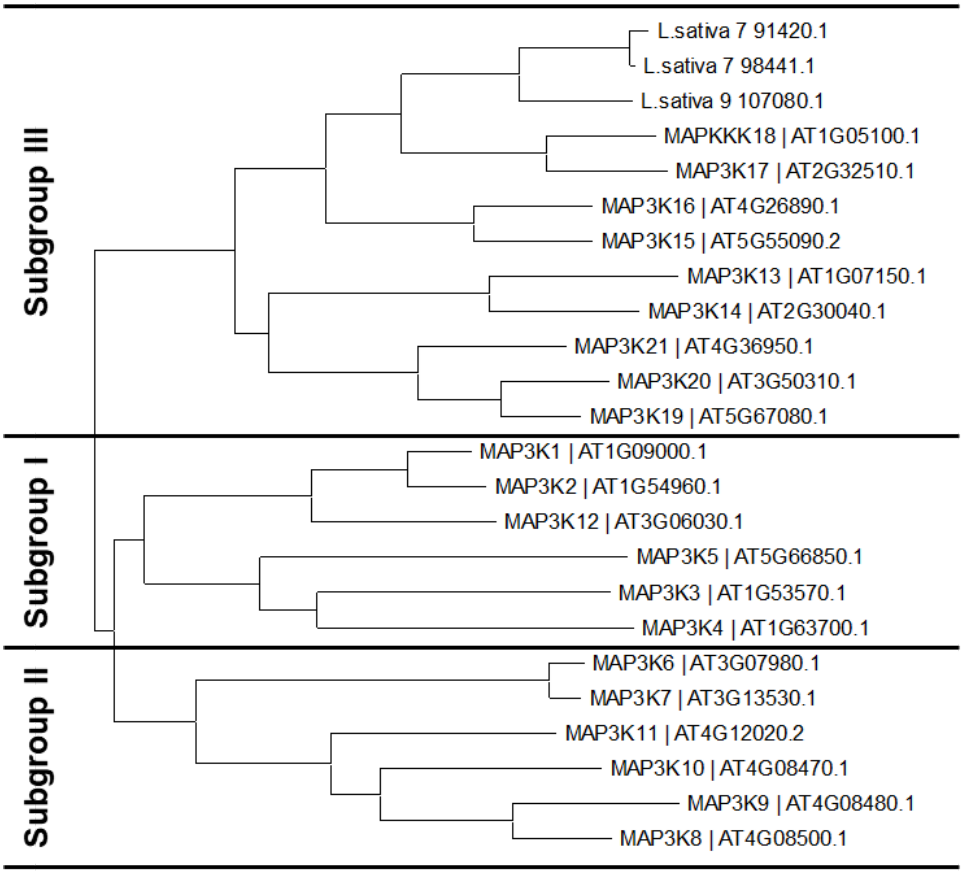


**Figure S1** Phylogenetic analysis of the family of Arabidopsis MEKK-like MAP3Ks including *Lactuca sativa* homologs. The tree shows the three subgroups of MAP3Ks in Arabidopsis and the clustering three lettuce homologs of MAP3K18 identified by BLAST.

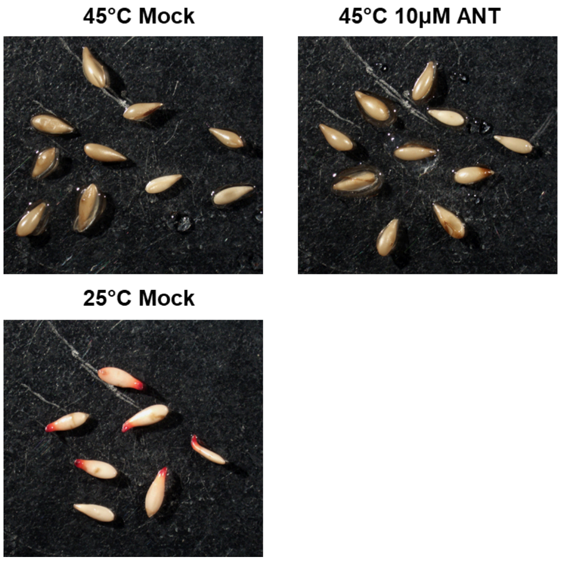


**Figure S2** *L. sativa* 'Salinas' seeds are no longer viable at 45°C. Seeds were imbibed in dark in 10μM ANT or mock treatment at 45°C for 120 hours before tetrazolium staining. These seeds were compared to mock treated seeds grown in dark for 24 hours before staining.

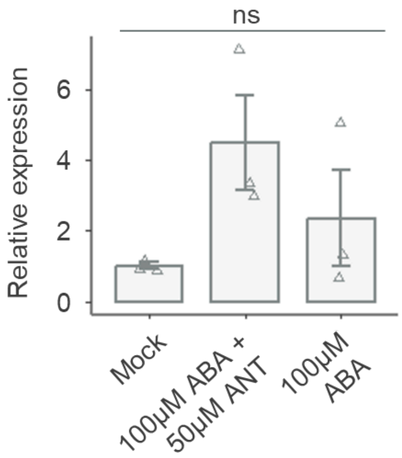


**Figure S3** ABA treatment does not significantly increase LsNCED4 expression. ABA and ABA + ANT treatments do not alter LsNCED4 expression (normalized to UBC21) measured by qRT-PCR of lettuce leaves in (DMSO) or 100μM ABA or 100μM ABA and 50μM ANT in water (pairwise t test, $p>0.05$)

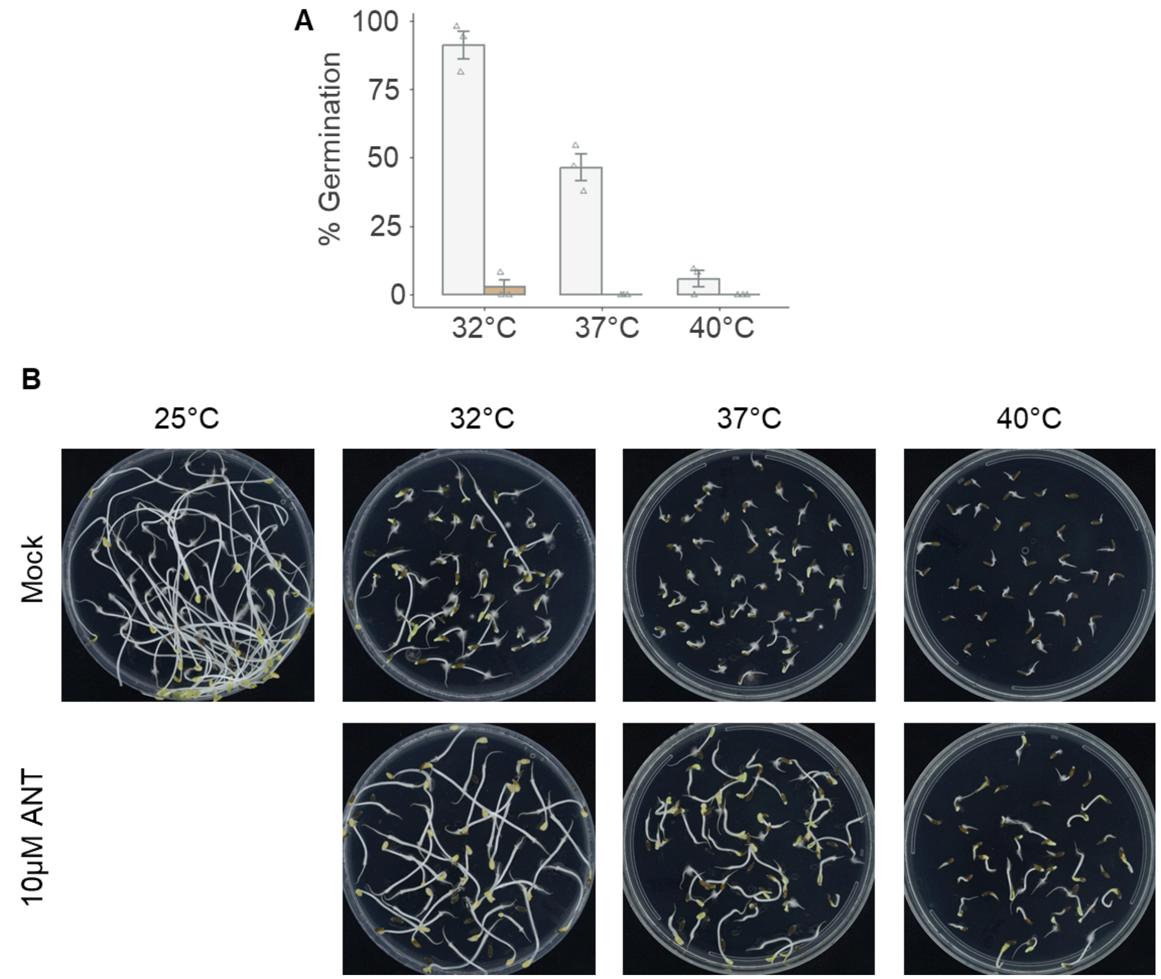


**Figure S4** ANT treatment restores germination at temperatures of up to 40°C. (A) Seeds treated with 10µM ANT begin to germinate at temperatures of up to 40°C after 48 hours of imbibition. (B) After 120 hours of heat treatment and 72 hours of recovery in dark at 25°C, seeds germinate.
